# Mechanism of physical force induced human slack channel desensitization

**DOI:** 10.1101/2023.01.07.523125

**Authors:** Mingfeng Zhang, Yuanyue Shan, Duanqing Pei

## Abstract

Human Slack (hSlack) channel is abundantly expressed in the central nervous system and regulates numerous physiological processes. Here, we identified hSlack channel is a voltage dependent physical force induced desensitization potassium channel. In the cell-attached mode, hSlack exhibited large voltage dependent potassium currents without significant delayed rectification or C-type inactivation. After breaking the cell membrane resulting the inside-out mode, the potassium currents were largely inhibited and it transited to the strong voltage dependent delayed rectified state. Structural results suggested that the “S6 site” confers the voltage dependent delayed rectification of hSlack channel, like our recent finding in hEag2 channel *(In revision)*. Strikingly, the physical force induced desensitization and gating kinetics transition are mediated by the gatekeeper lipids in the central pore. Our results reported a novel physical force sensing channel through the pore lipids mediating physical force induced desensitization and gating kinetics transition. It opens a new door to understand the molecular basis of mechanobiology.

## Introductions

The two human slowpoke (Slo) superfamily genes human Slack (Slo 2.2, KCNT1) and human Slick (Slo2.1, KCNT2) that are proposed to encode sodium activated potassium channel (K_Na_) channels^1,2^. Abnormal especially the gain of functions (GOF) of human Slack (hSlack) channel can result in severe early-onset epilepsy and neurodevelopmental disorders^3-5^. Therefore, the hSlack channel may server as a drug target to treat the early-onset epilepsy^5^. In animal models, the Slack channel regulates various physiological processes including nociceptive behavior^6^, itch^7^, neuropathic pain^8^, etc. Since these sensory inputs are physical factors, there is a question that whether slack channel can directly response to these physical cues.

The cryo-EM structures of chicken Slack (cSlack) channel illuminated the architectures of Slack channel. Additionally, the structures give evidences of conformational changes from the closed to open states through the inner helix of TM6 dilation^9^. However, the mechanisms of unique gating kinetics are still elusive due to the relatively low resolution and heterogeneities of the channel, including both carboxyl -terminal domain (CTD) and transmembrane domain (TMD).

In this study, we identified that hSlack channel is a voltage dependent physical force induced desensitization potassium channel and uncovered the molecular basis of force sensing via combining electrophysiological and structural studies. The physical force can inhibit the hSlack activity and force the channel into the delayed rectified state. Then, we solved two high resolution cryo-EM structures of human Slack channel at pre-open delayed rectification state, illuminating the delayed rectified process. Strikingly, four lipids in the central pore not only form the hydrophobic constrict site but also mediate the physical force induced desensitization and gating kinetics transition. Mutating the coordinated residues of the lipid results channel activation instead of inhibition when breaking the cell membrane. Meanwhile, the channel exhibits the C-type inactivation gating kinetics instead of delayed rectification. Additionally, the lipid binding site mutation acquire the mechanosensitive property, that can be activated by relatively high negative pressure. Overall, our results identified a novel physical force sensing ion channel and revealed the force sensing mechanism through the gatekeeper lipid. It will begin a feast to study this special kind of mechanobiology.

## Results

### human Slack channel is a physical force induced desensitization potassium channel

We expressed full-length hSlack channel fused with a green fluorescent protein (GFP) in HEK293T cells and determined its electrophysiological properties. As hSlack is a potassium channel, we used the asymmetric potassium solution to monitor the potassium current. Under cell-attached configuration without membrane disruption, the hSlack channel exhibited large voltage-dependent potassium currents without delayed rectification or C-type inactivation (Fig. 1a). Surprisingly, we reproducibly observed the fast inhibition manner while we pulled the membrane out of the cell forming the inside out configuration (Fig. 1b, g, i). Then, the hSlack channel showed the typical delayed rectified potassium currents with the activation time (τ) of 1046 ± 125 ms (Fig. 1c, k). Sodium is long thought the activator of Slack channel. To test the role of sodium in physical force induced desensitization and gating kinetics transition, we replaced the bath solution by 150 mM potassium without sodium and the pipette solution is 150 mM sodium, then performed electrophysiological study as before. The voltage dependent Slack channel currents were still inhibited when the cell membrane was disrupted by physical force (Extended Data Fig. 1c-e), suggesting that sodium do not participate in these processes. To further confirm the physical force induced desensitization phenotype, we tested the clinical screened GOF mutation F327L, which cause the early-onset epilepsy. The F327L also exhibited the physical force induced desensitization (Fig. 1d-f, h, j). Interestingly, the GOF mutation F327L attenuated the delayed rectification (Fig. 1k), indicating that the physical force induced gating kinetics transition may play an important role in the neuron system.

**Fig 1.**
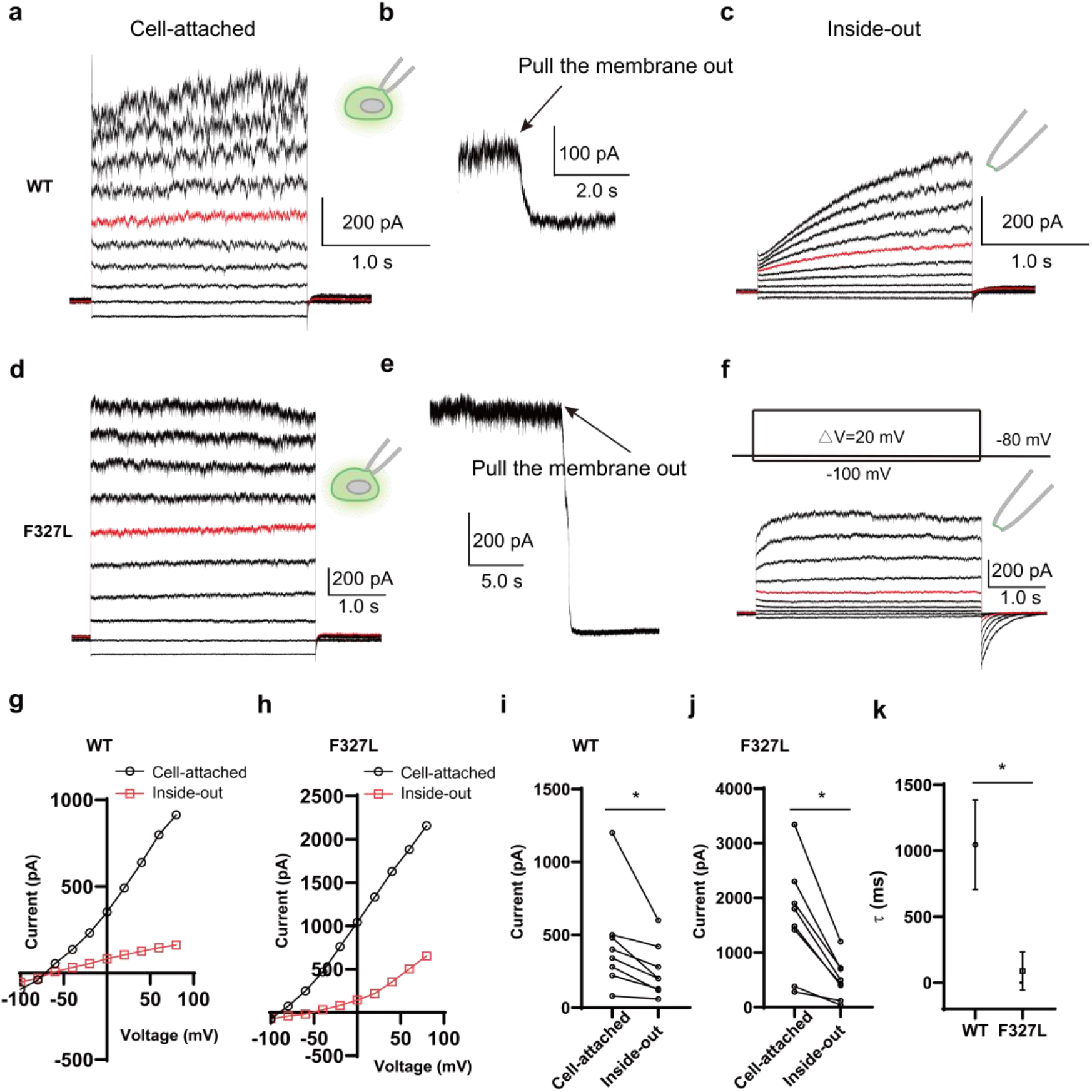
hSlack channel is a physical force induced desensitization potassium channel. **a**, Representative electrophysiological recording from the HEK293T cell expressing full-length wild-type hSlack with the voltage-pulse protocol from -100 mV to +80 mV with 20 mV per step under cell-attach mode. The bath solution was 10 mM HEPES-Na pH 7.4, 150 mM NaCl, 5 mM glucose, 2 mM MgCl_2_ and 1 mM CaCl_2_ and the pipette solution was 10 mM HEPES-Na pH 7.4, 150 mM KCl, 5 mM EGTA, therefore it generates a near physiological potassium gradient. The red trace stands for the current recorded at the holding potential of 0 mV. **b**, The current from the same cell as **(a)** at a holding potential of 0 mV. The arrow indicates pulling the membrane from the cell and forming the inside-out model. **c**, Representative electrophysiological recording from the same cell as **(b)** with the voltage-pulse protocol from -100 mV to +80 mV with 20 mV per step after forming inside-out mode. **d-f**, Representative electrophysiological recording from the HEK293T cell expressing hSlack-F327L mutation. The recording modes are corresponding to **(a-c). g-h**, I-V plot of wildtype hSlack **(g)** and hSlack-F327L mutation **(h)** under different recording modes from **(a, c)** and **(d, f)** respectively.

### Structure determination of human Slack channel

To study the mechanism of physical force induced desensitization and gating kinetics transition, we purified the full length hSlack channel in a 150 mM KCl environment and subjected it to the standard cryo-EM workflow. After cryo-EM imaging and data processing (Extended Data Fig. 2), we isolated two distinct near-atomic resolution high quality maps at overall resolution of 3.3 (PDB:8HLI; EMD-34877), and 3.2 Å (PDB:8HLH; EMD-34876) respectively (Extended Data Fig. 2-5). The map has a distinct two-layer architecture: transmembrane domains (TMD) and cytosolic domain (CTD) (Fig. 2a-f). One map exhibits high quality of TMD (Fig. 2a-c) and another shows high quality of CTD and the pore region (Fig. 2d-f). These two high-quality maps allowed us to build an accurate near-atomic model of hSlack at different conformations (Extended Data Fig. 4, Extended Data Table. 1). Each hSlack subunit contains N-terminal domain, voltage-sensor domain (VSD) formed from the S1-S4 transmembrane helices, pore domain formed from the S5-pore helix (PH)-S6 domain and the C-terminal RCK1 and RCK2 domain (Fig. 2g-h). The overall tetrameric structure of hSlack is identical to the cSlack^9^ and calcium activated BK^10^ channels with the non-domain-swapped conformation between pore domain and VSD, but the domain-swapped conformation between TMD and CTD (Fig. 2i-j).

**Fig 2.**
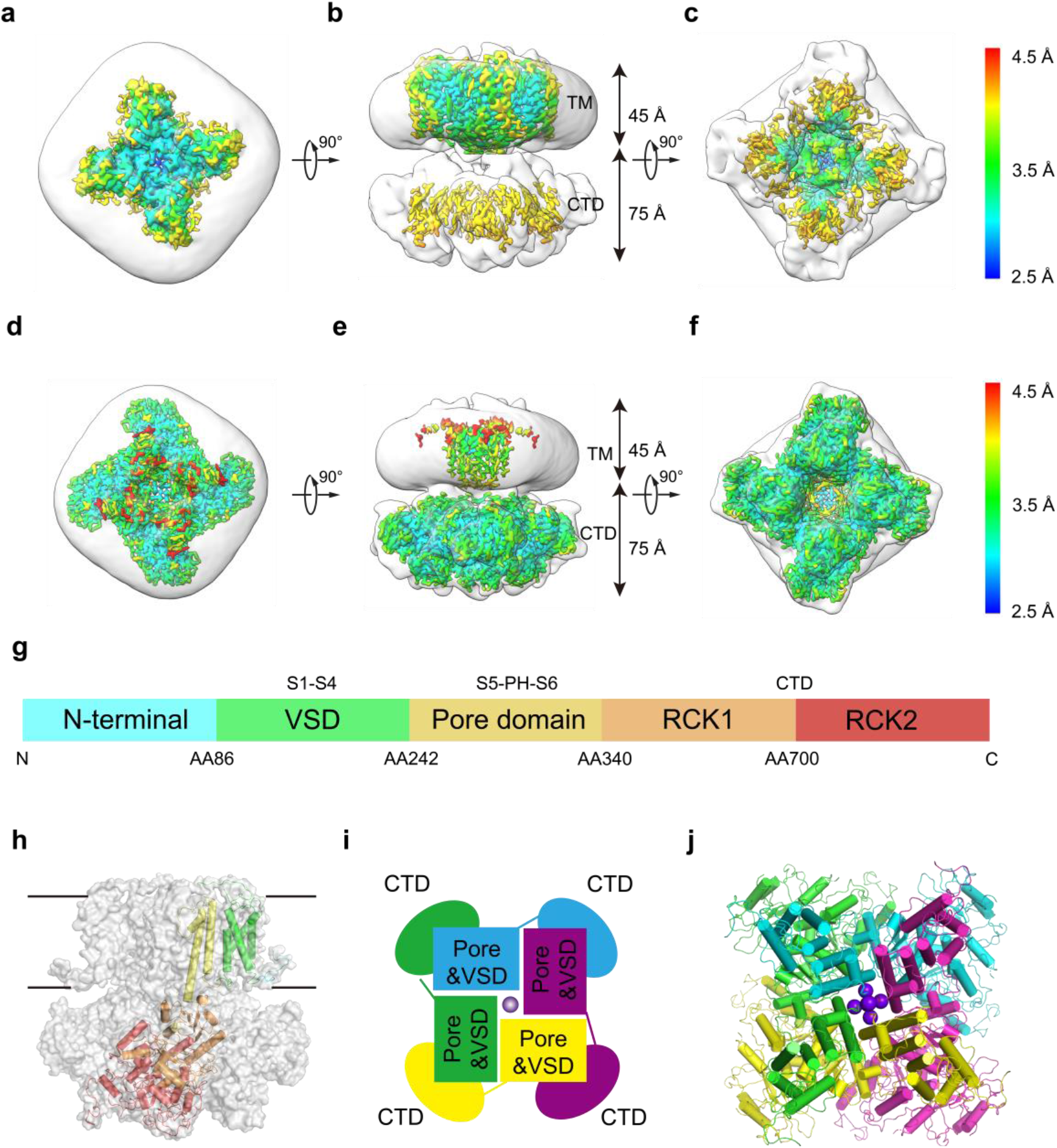
Overall structure of hSlack channel. **a-c**, Top view **(a)**, side view **(b)** and bottom view **(c)** of TMD high quality cryo-EM density map with detergent micelle (grey) and estimated resolution are shown. **d-f**, Top view **(d)**, side view **(e)** and bottom view **(f)** of CTD high quality cryo-EM density map with detergent micelle (grey) and estimated resolution are shown. **g**, Schematic representation of hSlack domain structure. Domains are rendered in colors according to the protomer structure shown in **(h). h**, One subunit is displayed as cartoon and the remaining subunits are shown in surface representation. **i**, Carton presents the non-domain-swapped between TMD and CTD, TMD (pore domain and VSD) and CTD of hEag2 tetramers in top view. A potassium ion in the pore domain is depicted as a purple sphere. **j**, Overall structure of hSlack tetramer in top view. Each subunit is rendered in colors.

### The pre-open state 1 and pre-open state 2 of hSlack

The closed and open state of cSlack model reveals the inner helix of TM6 dilation during channel activation^9^. The pore sizes of two model are closed to the closed state of cSlack channel, therefore it is at least not the fully open state (Fig. 3a-f). Our previous work reveals the delayed rectification mechanism of huma Eag2 (hEag2) channel, suggesting that the constrict sites of S6 (formed by T468s and Q472s) keep “closed” confer the delayed rectification upon the voltage gradient stimuli. In hSlack channel, the corresponding the constrict sites of S6 are formed by the M335s and Q338s. In the TMD high quality model, one potassium ion occupies in the constrict site of M335s and waters occupy in the constrict site of Q338s (Fig. 3g) while in the CTD high quality model, one potassium ion occupied in the constrict site of M335s and a pre-dehydrated potassium (one potassium ion is surrounded by waters) occupied in the constrict site of Q338s (Fig. 3h). We name the TMD high quality map as pre-open state 1 and CTD high quality map as pre-open state 2. The potassium dynamic process in the constrict sites of S6 site give us a clear picture of the potassium ion dehydration process (Fig. 4a-b). Therefore, we proposed that the two pre-open states stand for the delayed rectified states of hSlack channel, which consistent with hSlack channel extracting from the cell membrane like the inside-out mode. In the hEag2, the Arginine replacement of glutamine at “S6” eliminate the delayed rectified property. We mutated the Q338 to arginine and then test the electrophysiological data as before. Interestingly, the delayed rectified property of Q338R is instead by the typical C-type inactivation but remain the physical force induced desensitization (Fig. 4c-e).

**Fig 3.**
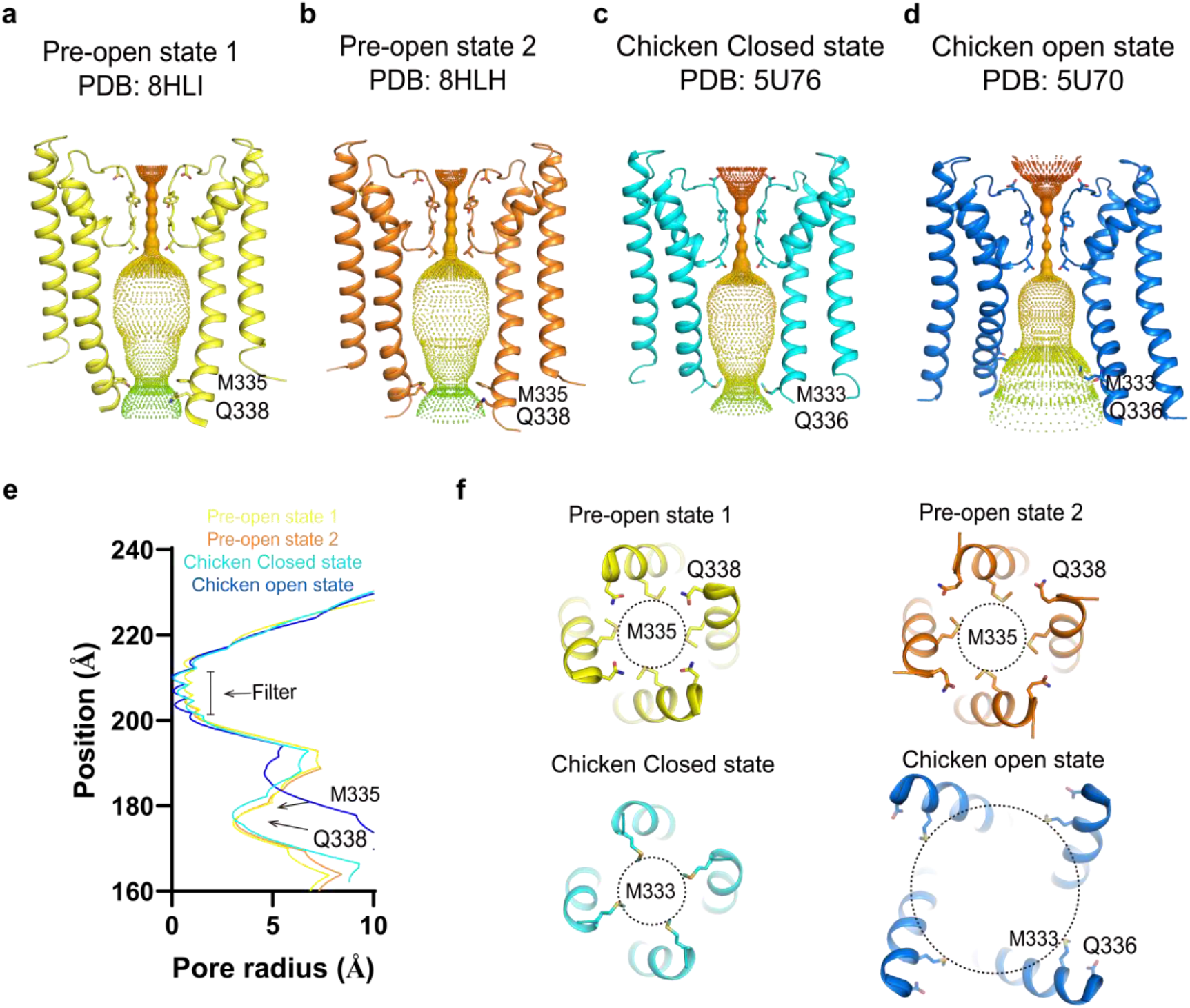
Structural comparison of human and chicken Slack channel pore domain. **a-d**, Ion pore with only two subunits of hSlack in pre-open state 1 (**a**); hSlack in pre-open state 2 (**b**); chicken Slack in closed state (**c**); chicken slack in open state (d) shown, viewed from within the membrane. The minimal radial distance from the center axis to the protein surface is colored in iridescence. Selected residues facing the pore are in stick representation. **e**, Comparison of the pore radius of hSlack in pre-open state 1, hSlack in pre-open state 2, chicken Slack in closed state and chicken slack in open state. The van der Waals radius is plotted against the distance along the pore axis. The selectivity filter and the constricting residues are labeled. **f**, Enlarged top view of the pore domain constricting residues in hSlack (M335 and Q338) and chicken Slack (M333 and Q336). The size of the pore is present as black dashed circle.

**Fig 4.**
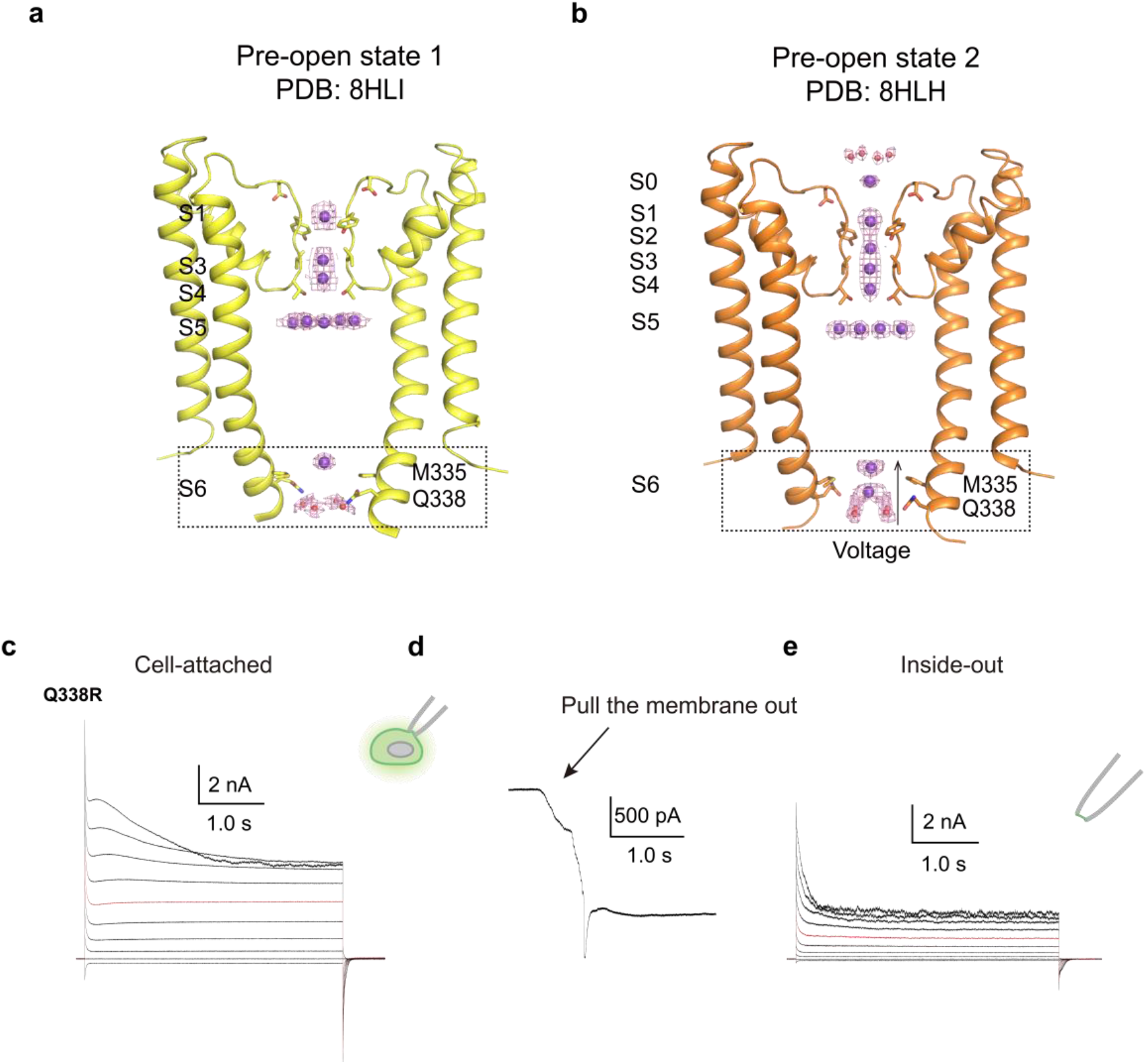
The pre-open state 1 and pre-open state 2 of hSlack. **a-b**, Selectivity filter and central cavity of hSlack in pre-open state 1 (**a**) and pre-open state 2 (**b**) shown, viewed from within the membrane. Water molecules above the selectivity filter are shown as red spheres and the potassium ions in S0 to S6 are shown as purple spheres with electron density maps (purple mesh). The selectivity residues and pore domain constricting residues are represented as sticks. **a**, Representative electrophysiological recording from the HEK293T cell expressing Q338R with the voltage-pulse protocol from -100 mV to +80 mV with 20 mV per step under cell-attach mode. The bath solution was 10 mM HEPES-Na pH 7.4, 150 mM NaCl, 5 mM glucose, 2 mM MgCl_2_ and 1 mM CaCl_2_ and the pipette solution was 10 mM HEPES-Na pH 7.4, 150 mM KCl, 5 mM EGTA, therefore it generates a near physiological potassium gradient. The red trace stands for the current recorded at the holding potential of 0 mV. **b**, The current from the same cell as **(a)** at a holding potential of 0 mV. The arrow indicates pulling the membrane from the cell and forming the inside-out model. **c**, Representative electrophysiological recording from the same cell as **(b)** with the voltage-pulse protocol from -100 mV to +80 mV with 20 mV per step after forming inside-out mode.

### Lipids occupied in the central pore forming a hydrophobic constrict site

Since the two models stand for the delayed rectified process of hSlack channel under inside-out mode, some molecular will mediate the physical force induced desensitization. One of most promising candidates is lipid. Interestingly, four clear lipid densities (each subunit contains one) are occupied in the central pore below the selectivity filter (Fig. 5a-d). We call these lipids as gatekeeper lipids. The gatekeeper lipids together with the L320s on the TM6 form a hydrophobic constrict site. At least five clear ion densities (they most probably are potassium ions or waters) occupy in the hydrophobic constrict site like a potassium ion “reservoir” (Fig. 5g-e). In hEag2, there is also a hydrophobic constrict site upon the constrict sites of S6. Despite the constrict sites of S6 keep closed in the multiple sub-fully open state, the hydrophobic constrict site undergo multiple conformational changes to assist the ion flow under the delayed rectified state. To test the role of hydrophobic constrict site in hSlack channel gating, we mutated the L320 to alanine and found that the single channel conductance of the mutation is largely reduced (Fig. 5f), suggesting it functions an important role in ion flow rate and support the function of potassium ion “reservoir”.

**Fig 5.**
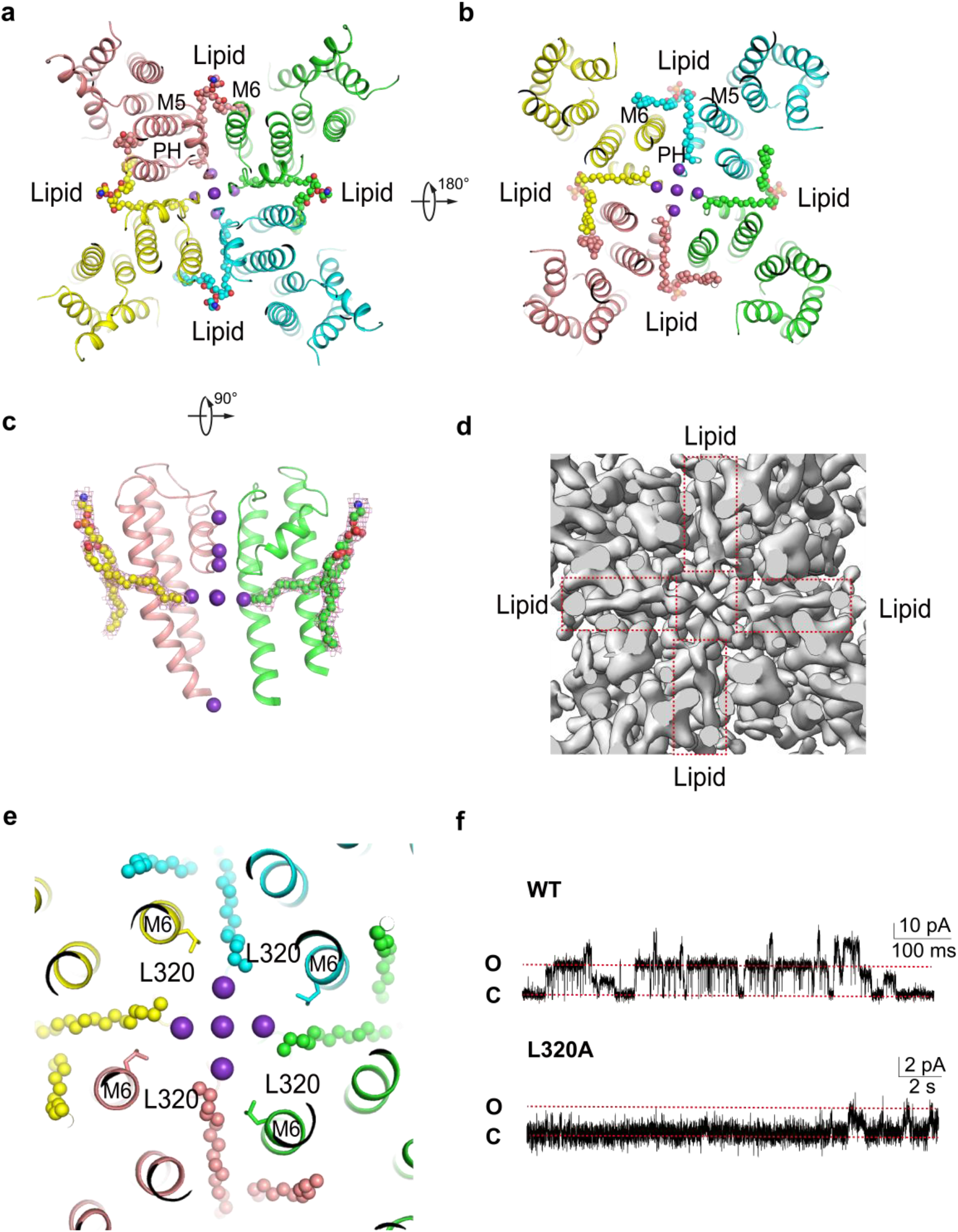
Lipids occupied in the central pore forming a hydrophobic constrict site. **a-c**, Lipid binding site of hSlack in closed state viewed from top (**a**), bottom (**b**) and side (**c**). The four subunits are shown as cartoon in different colors and the lipids are shown as spheres. **d**, Density map of lipids binding site of hSlack in closed state. The lipids density is highlighted by red dash boxes. **e**, Key residues L320 in lipids biding site on M6, viewed from the top. **f**, Representative single-channel currents of wild-type hSlack and L320A mutant at a holding potential of 0 mV.

### The gatekeeper lipids mediate physical force induced desensitization and gating kinetics transition

The gatekeeper lipids not only form the hydrophobic constrict site, but also interact with the TM5, TM6 and the pore helix (PH) (Fig. 6a-b). To further investigate the potential role of gatekeeper lipids in channel gating, we mutated the residues which directly interact with the gatekeeper lipids. The interaction mutation I290A at the PH domain display relative silent under the cell attach mode (Fig. 6c). Strikingly, when forming the inside-out mode, the channel is largely activated instead of inhibited (Fig. 6d). Also, the mutation I290A channel showed the typical fast C-type inactivation currents (Fig. 6e-f). Now that the I290A can be activated by the membrane disruption, we suspected it can be activated by membrane stretch. We applied the negative pressure to the membrane under cell-attached mode at the holding potential of 0 mV to recorded the typical mechanosensitive potassium currents. Owing to the hSlack channel is the potassium channel, it is clear to distinguish the noise or the potential innate weak background piezo1 current at the holding potential of 0 mV. The I290A mutation display the typical mechanosensitive current under cell-attached mode at the holding potential of 0 mV (Fig. 6g). Additionally, we found the V263A mutation on the TM6 interacting I290 and the gatekeeper lipids is also activated by the negative pressure, displaying well mechanosensitive property (Fig. 6h). Therefore, these results strongly indicate that the gatekeeper lipids mediate the physical force induced desensitization and gating kinetics transition, which also obey the classical “force-from-lipid” principle^11^.

**Fig 6.**
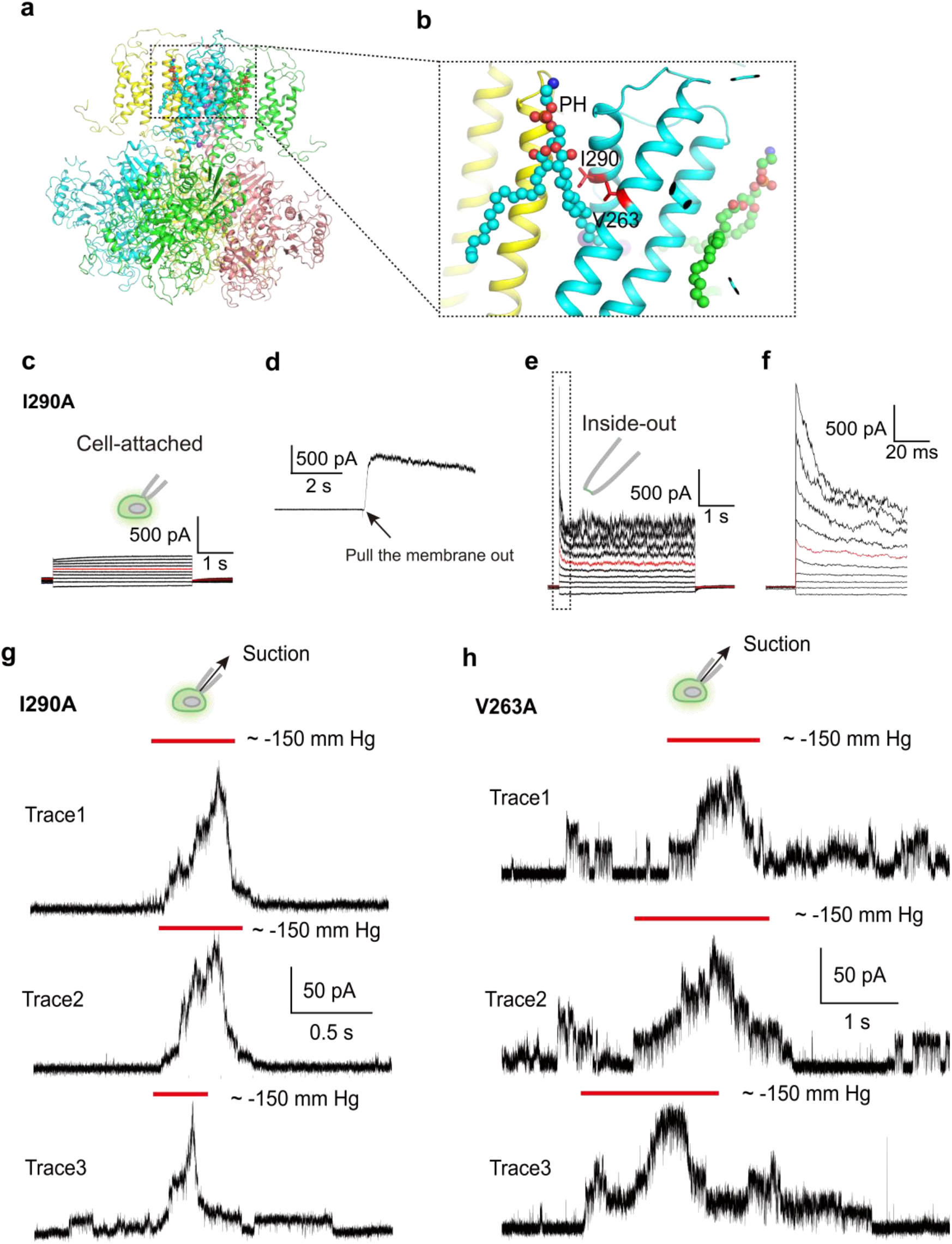
The gatekeeper lipids mediate physical force induced desensitization and gating kinetics transition. **a-b**, lipid binding site in pre-open 1 state hSlack (**a**) and enlarged lipid binding site (**b**) are shown in the side view. Residues interacting with lipids on pore helices (PH) are labeled. **c**, Representative electrophysiological recording from the HEK293T cell expressing hSlack mutant I290A with the voltage-pulse protocol from -100 mV to +80 mV with 20 mV per step under cell-attach mode. The red trace stands for the current recorded at the holding potential of 0 mV. **d**, The current from the same cell as (**c**) at a holding potential of 0 mV. The arrow indicates pulling the membrane from the cell and forming the inside-out model. **e**, Representative electrophysiological recording from the same cell as (**d**) with the voltage-pulse protocol from -100 mV to +80 mV with 20 mV per step after forming inside-out mode. **f**, Enlarged view of black dashed box in **e. g-f**, Representative traces of negative-pressure-activated currents recorded from cells transfected with hSlack mutants I290A and V263A in cell-attached mode at a holding potential of 0 mV. Negative pressure was applied around -150 mm Hg.

## Discussion

### Mechanism of delayed rectification of hSlack channel

The channel gating kinetics including C-type/ N-type inactivation^12-14^, delayed rectification^15^ etc. al fine tune the cellular ion homeostasis. Compare with the C-type/ N-type inactivation, the mechanism of delayed rectification is much younger. Our recent work illuminated the atomic mechanism of the typical voltage gated delayed rectified potassium channel human Eag2. During channel activation, the channel undergoes multiple sub-fully open states, while the constrict site (S6) of multiple sub-fully open states do not extend. We hypothesized that the S6 may conduct the ion upon the voltage acting on it. Our two pre-open state hSlack map clear present the potassium dehydration process, which supporting our hypotheses of the S6 simply senses the voltage gradient and then dehydration of water thus mediate channel delayed-rectification.

### Mechanism of physical force sensing of hSlack

Cells sense the external environment physical force changes incessantly^16-18^. The mechanosensitive ion channels play the major roles in the physical force sensing through regulating the cellular ion homeostasis^19^. In eukaryotic kingdom, the well-known so-called bona-fide mechanosensitive ion channels are only Piezo^20^, OSCA^21,22^ and K2P^23^ channels. Despite the multiplex 3D architectures of these channels^21,24-26^, they obey the basic force-from-lipid principle^11^. Also, all of them can be activated by membrane stretch despite they have different pressure threshold^20,21^. Our results demonstrate that the wildtype hSlack channel also response the physical force and obey the force-from-lipid principle despite the way of sensing physical force is quite different from the Piezo, OSCA and K2P channel. However, there also seems a conserved principle of gatekeeper lipid occupied in the pore^26,27^. Whether the gatekeeper lipid is the bone-fide principle remains more structural results to uncover them.

Limited to the electrophysiology method, the higher pressure may disrupt the membrane. It will fail to form the giga-seal and cannot record the signal. However, as the cells especially the neurons may expose to many complicated physical force environments. For example, the cilia, dendrites, etc. have unique cellular structures may challenge high physical force range. Meanwhile, some special physiological process such as cell division, cell migration, vesicle releasing, etc. may generate quiet high pressure to the cells. Therefore, we suspected that hSlack may play important functions in these situations. Further in vivo physiological roles of hSlack channel are waiting to be discovered.

## Materials and Methods

### Construct design

DNA fragment encoding hSlack channel (UniProtKB Q6UVM3) were synthesized (Tsingke Biotechnology) and cloned into the plasmid Eric Gouaux (pEG) BacMam vector using EcoRI and XhoI restriction sites, where hSlack gene is placed before a PreScission protease (Ppase) cleavage site, followed by a C-terminal enhanced green fluorescent protein (EGFP) and FLAG tag. The construct was identified by fluorescence detection size-exclusion chromatography (FSEC)^28^.

### Protein Expression and purification

The hSlack was expressed in expi-HEK293F cells using the BacMam method. Briefly, a bacmid carrying hSlack was generated by transforming E. *coli* DH10Bac cells with the hSlack construct according to the manufacturer’s instructions (Bac-to-Bac; Invitrogen). Baculoviruses were produced by transfecting Sf9 cells with the bacmid using Cellfectin II (Invitrogen). After three rounds of amplification, baculoviruses were used for cell transduction. Suspension cultures of expi-HEK293F cells were grown at 37°C to a density of ∼3×10^6^ cells/ml and viruses were added (8% v/v) to initiate the transduction. After 12 h, 10 mM sodium butyrate was supplemented, and the temperature was shifted to 30°C. Cells were harvested at 60 h post-transduction.

The cell pellet from 6 L culture was collected by centrifugation at 4,000 r.p.m. and resuspended with 500 ml of lysis buffer (20 mM Tris-HCl, pH 8.0, 150 mM KCl), supplied with protease inhibitor cocktail containing 1.3 μg/ml aprotinin, 0.7 μg/ml pepstatin, and 5 μg/ml leupeptin and 2 mM phenylmethylsulfonyl fluoride (PMSF). Unbroken cells and cell debris were removed by centrifugation at 8,000 r.p.m. for 10 min. Supernatant was centrifuged at 36,000 r.p.m. for 30 min in a Ti45 rotor (Beckman). The membrane pellet was mechanically homogenized and solubilized in extraction buffer containing 20 mM Tris-HCl, pH 8.0, 150 mM KCl, 1% (w/v) lauryl maltose neopentyl glycol (LMNG) and 0.1% (w/v) cholesteryl hemisuccinate (CHS). Unsolubilized materials were removed by centrifugation at 36,000 r.p.m. for 1 h in a Ti45 rotor. Supernatant was load onto anti-FLAG G1 affinity resin (Genscript) by gravity flow. Resin was further washed with 10 column volumes of wash buffer (lysis buffer with 0.02% LMNG), and protein was eluted with an elution buffer (lysis buffer with 0.02% LMNG and 230 μg/ml FLAG peptide). The C-terminal GFP tag of eluted protein was removed by human rhinovirus 3C (HRV 3C) protease cleavage for 2 h. The protein was further concentrated by a 100-kDa cutoff concentrator (Milipore) and loaded onto a Superose 6 increase 10/300 column (GE Healthcare) running in lysis buffer with 0.03% digitonin. Peak fractions were pooled and concentrated to around 5 mg/ml for cryo-EM sample preparation. All steps were performed at 4°C.

### Cryo-EM sample preparation

For cryo-EM sample preparation, aliquots (3.5 μl) of the protein sample were loaded onto glow-discharged (20 s, 15 mA; Pelco easiGlow, Ted Pella) Au grids (Quantifoil, R1.2/1.3, 300 mesh). The grids were blotted for 3.5 s with 3 force after waiting for 5 s and immersed in liquid ethane using Vitrobot (Mark IV, Thermo Fisher Scientific/FEI) in condition of 100% humidity and 8°C.

### Cryo-EM image acquisition

Cryo-EM data for hSlack were collected on a Titan Krios microscope (FEI) equipped with a cesium corrector operated at 300 kV. Movie stacks were automatically acquired with EPU software^29^ on a Gatan K3 Summit detector in super-resolution mode (105,000× magnification) with pixel size 0.4245 Å at the object plane and with defocus ranging from -1.5 μm to -2.0 μm and GIF Quantum energy filter with a 20 eV slit width. The does rate on the sample was 23.438 e^-^ s^-1^ Å^-2^, and each stack was 2.56 s long and does-fractioned into 32 frames with 80 ms for each frame. Total exposure was 60 e^-^ Å^-2^. Acquisition parameters are summarized in table S1.

### Image processing

A flowchart of the Cryo-EM data processing process can be found in Extended Data Fig. 1. Data processing was carried out with the cryoSPARC v3 suite^30^. Super-resolution image stacks were gain-normalized, binned by 2 with Fourier cropping, and patch-based CTF parameters of the dose-weighted micrographs (0.849 Å per pixel) were determined by cryoSPARC. Around 375K particles were picked by blob picking from 999 micrographs and 2D classification was performed to select 2D classes as templates. Template picker was used to pick around 1,351K particles. Particles were extracted using a 400-pixel box with a pixel size of 0.849 Å. The dataset was cleared using several rounds of 2D classification, followed by Ab initio reconstruction using C1 symmetry and initial model (PDB: 5U70). Two classes with 344.9K particles (34.7%) and 396.6 K particles (39.9%) were further refined using non-uniform refinement and local refinement for reconstructing the density map while imposing a C4 symmetry. The two final maps were low-pass filtered to the map-model FSC value. The reported resolutions were based on the FSC=0.143 criterion.

### Model building

The atomic models of monomer were built in Coot based on an initial model (PDB: 5U70). The models were then manually adjusted in Coot. Tetramer models were obtained subsequently by applying a symmetry operation on the monomer. These tetramer models were refined using phenix.real_space_refine with secondary structure restraints and Coot iteratively. Residues whose side chains have poor density were modeled as alanines. For validation, FSC curves were calculated between the final models and EM maps. The pore radii were calculated using HOLE. Figures were prepared using PyMOL and Chimera.

### Electrophysiological recording

HEK293T cells were cultured on coverslips placed in a 12-well plate containing DMEM medium (Gibco) supplemented with 10% fetal bovine serum (FBS). The cells in each well were transiently transfected with 1 μg hSlack DNA plasmid using Lipofectamine 3000 (Invitrogen) according to the manufacturer’s instructions. After 24-48 h, the coverslips were transferred to a recording chamber containing the external solution (10 mM HEPES-Na pH 7.4, 150 mM NaCl, 5 mM glucose, 2 mM MgCl_2_ and 1 mM CaCl_2_). Borosilicate micropipettes (OD 1.5 mm, ID 0.86 mm, Sutter) were pulled and fire polished to 2-5 MΩ resistance. The pipette solution was 10 mM HEPES-Na pH 7.4, 150 mM KCl, 5 mM EGTA. Both cell-attached and inside-out recordings were obtained at room temperature (∼25°C) using an Axopatch 700B amplifier, a Digidata 1550 digitizer and pCLAMP software (Molecular Devices). The patches were held at - 80 mV and the recordings were low-pass filtered at 1 kHz and sampled at 20 kHz. Statistical analyses were performed using Graphpad Prism.

**Extended Data Fig. 1.**
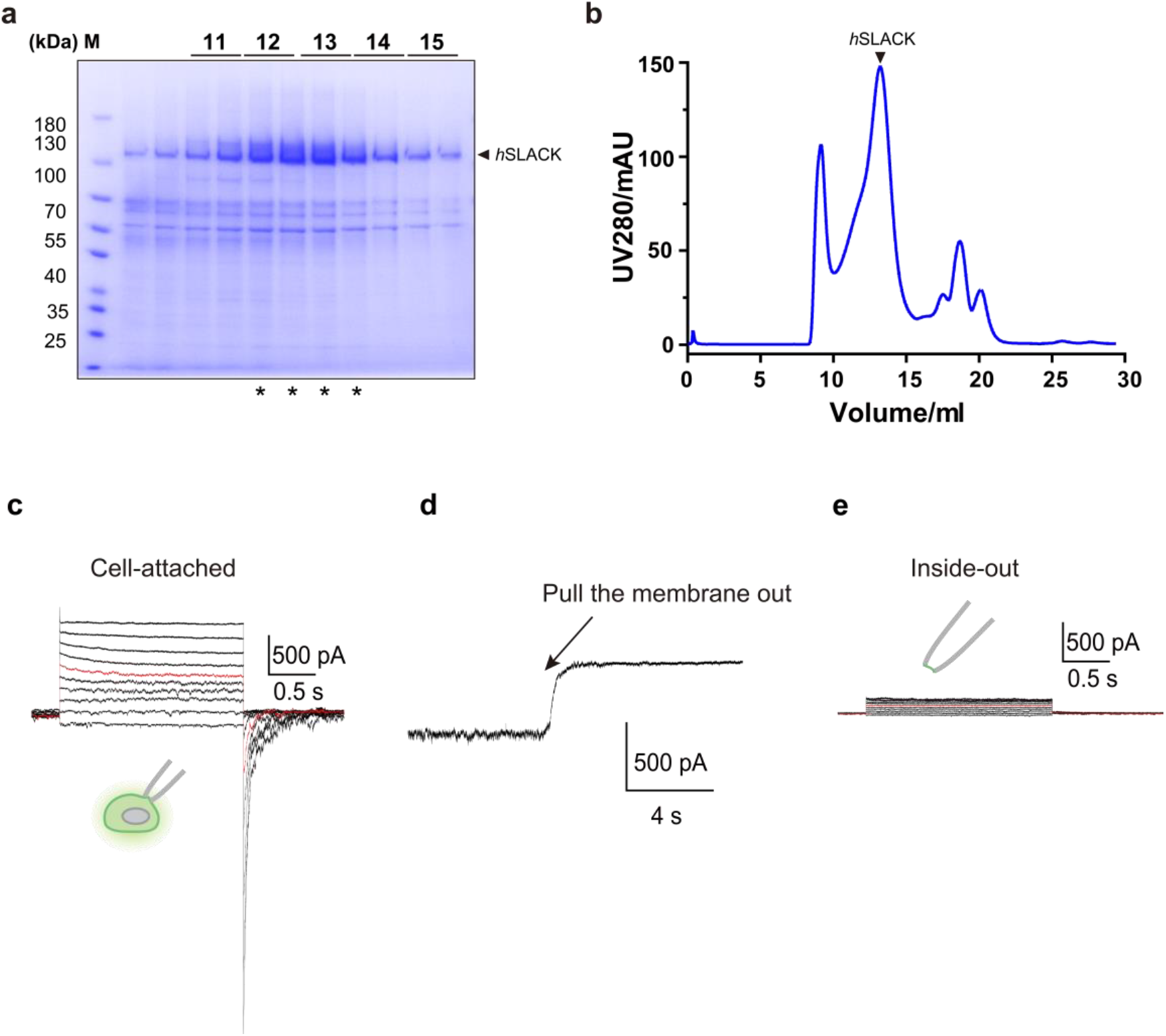
Purification of the hSlack channel. **a**, Representative size-exclusion chromatography (SEC) trace of purified hSlack channel. **b**, SDS-PAGE gel of peak fractions stained with Coomassie blue. The predicted molecular weight based on protein sequence for a single hSlack subunit is 138.3 kDa. Peak fractions labeled with stars were used for cryo-EM analysis. **c**, Representative electrophysiological recording from the HEK293T cell expressing full-length wild-type hSlack with the voltage-pulse protocol from -100 mV to +80 mV with 20 mV per step under cell-attach mode. The bath solution was 10 mM HEPES-K pH 7.4, 150 mM KCl, 5 mM EGTA and the pipette solution was 10 mM HEPES-Na pH 7.4, 150 mM NaCl, 5 mM glucose, 2 mM MgCl_2_ and 1 mM CaCl_2_. The red trace stands for the current recorded at the holding potential of 0 mV. **d**, The current from the same cell as **(c)** at a holding potential of 0 mV. The arrow indicates pulling the membrane from the cell and forming the inside-out mode. **e**, Representative electrophysiological recording from the same cell as **(d)** with the voltage-pulse protocol from -100 mV to +80 mV with 20 mV per step after forming inside-out mode.

**Extended Data Fig. 2.**
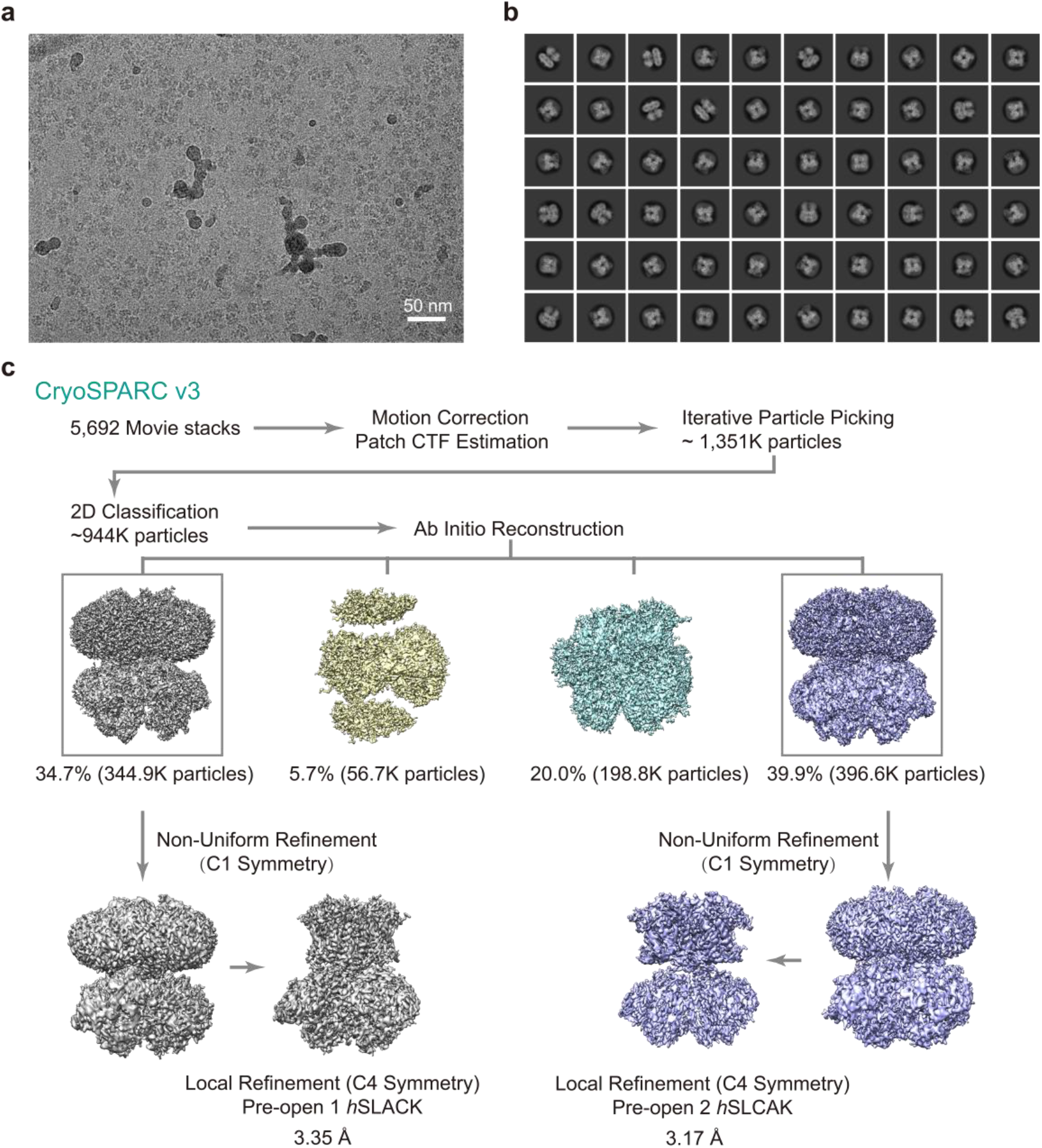
Single-particle Cryo-EM reconstructions of the hSlack channel. **a**, A representative raw micrograph of the hSlack. **b**, Selected 2D class averages. **c**, Summary of image processing for two states of hSlack dataset with C4 symmetry.

**Extended Data Fig. 3.**
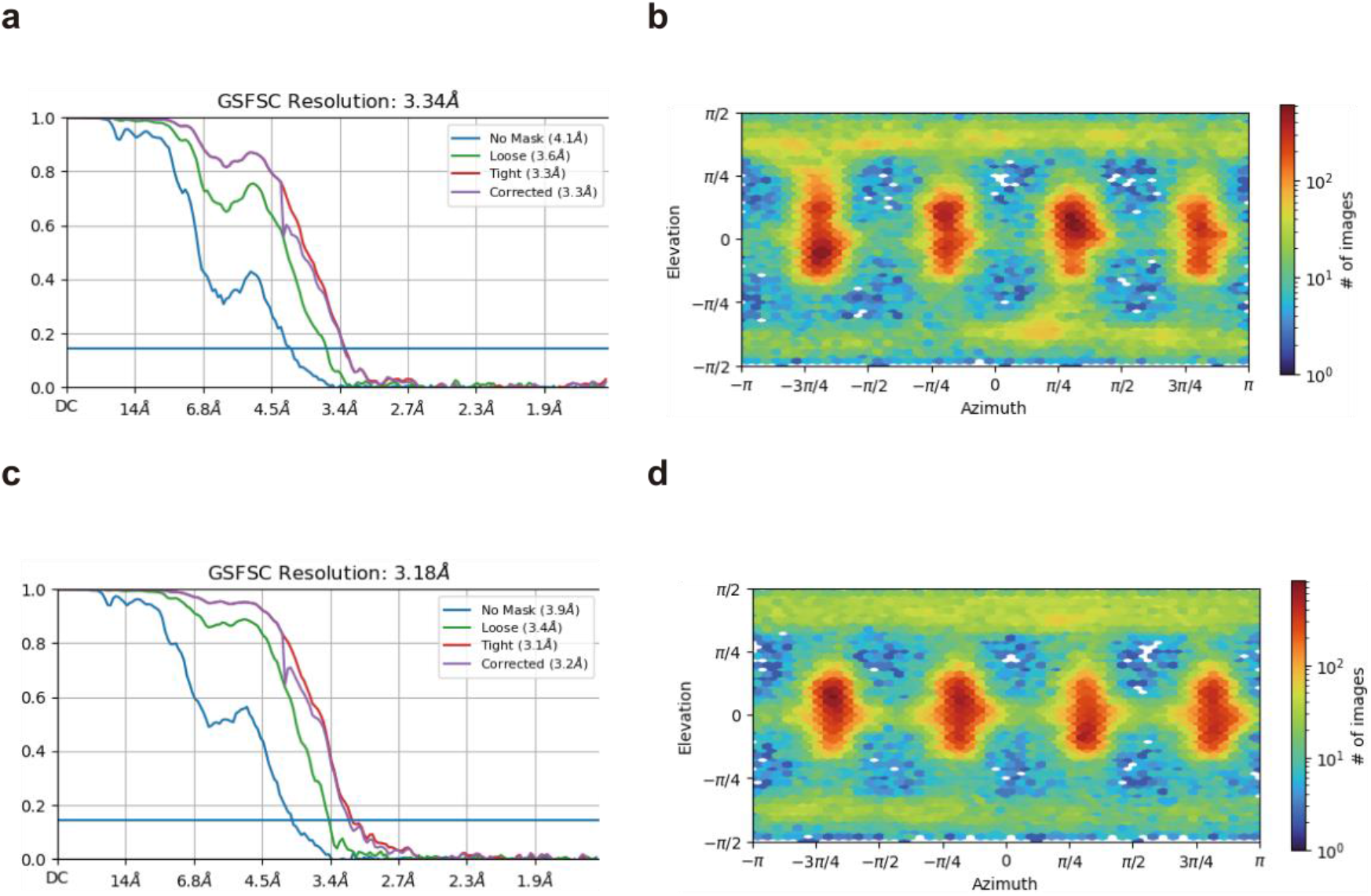
Fourier Shell correlation (FSC) curves and Euler angle distribution of particles for the final 3D reconstruction of hSlack in six states. FSC curves between two half maps before (blue) and after loose (Green) and tight (Purple) masks of hSlack in pre-open state 1 **(a)** and pre-open state 2 **(c)**, respectively. Euler angle distribution of particles for 3D reconstruction of hSlack in pre-open state 1 **(b)** and pre-open state 2 **(d)**, respectively. The reported resolutions were based on the FSC=0.143 criterion.

**Extended Data Fig. 4.**
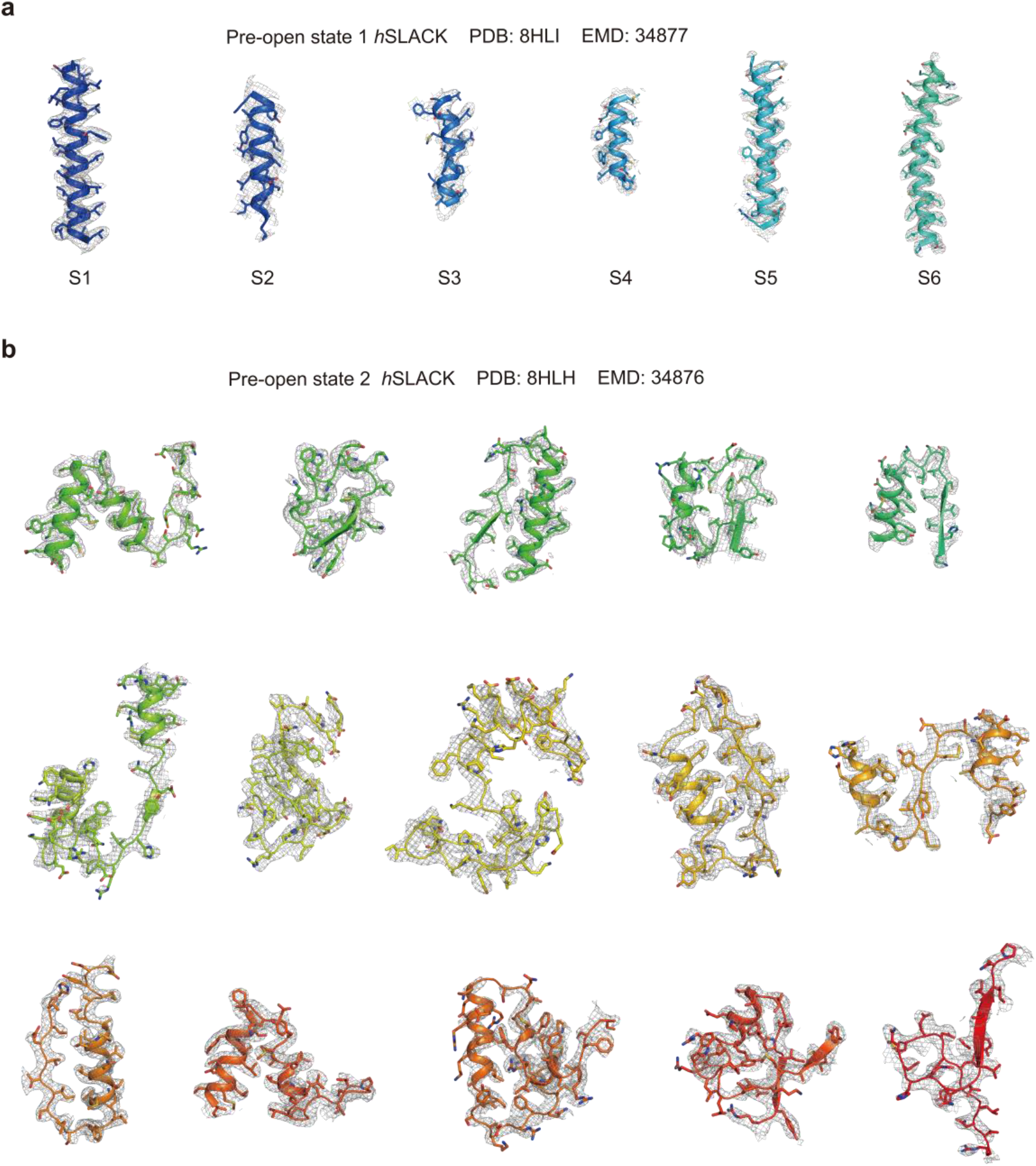
Local resolution of two hSlack density maps estimated by local resolution estimation in CryoSPARC. Local resolution of hSlack in pre-open state 1 **(a)** and pre-open state 2 **(b)** in bottom view (left), side view (middle) and top view (right). FSC curves for cross-validation: model versus unmasked full map (blue) and masked full map (purple) of hSlack in pre-open state 1 **(c)** and pre-open state 2 **(d)**.

**Extended Data Fig. 5.**
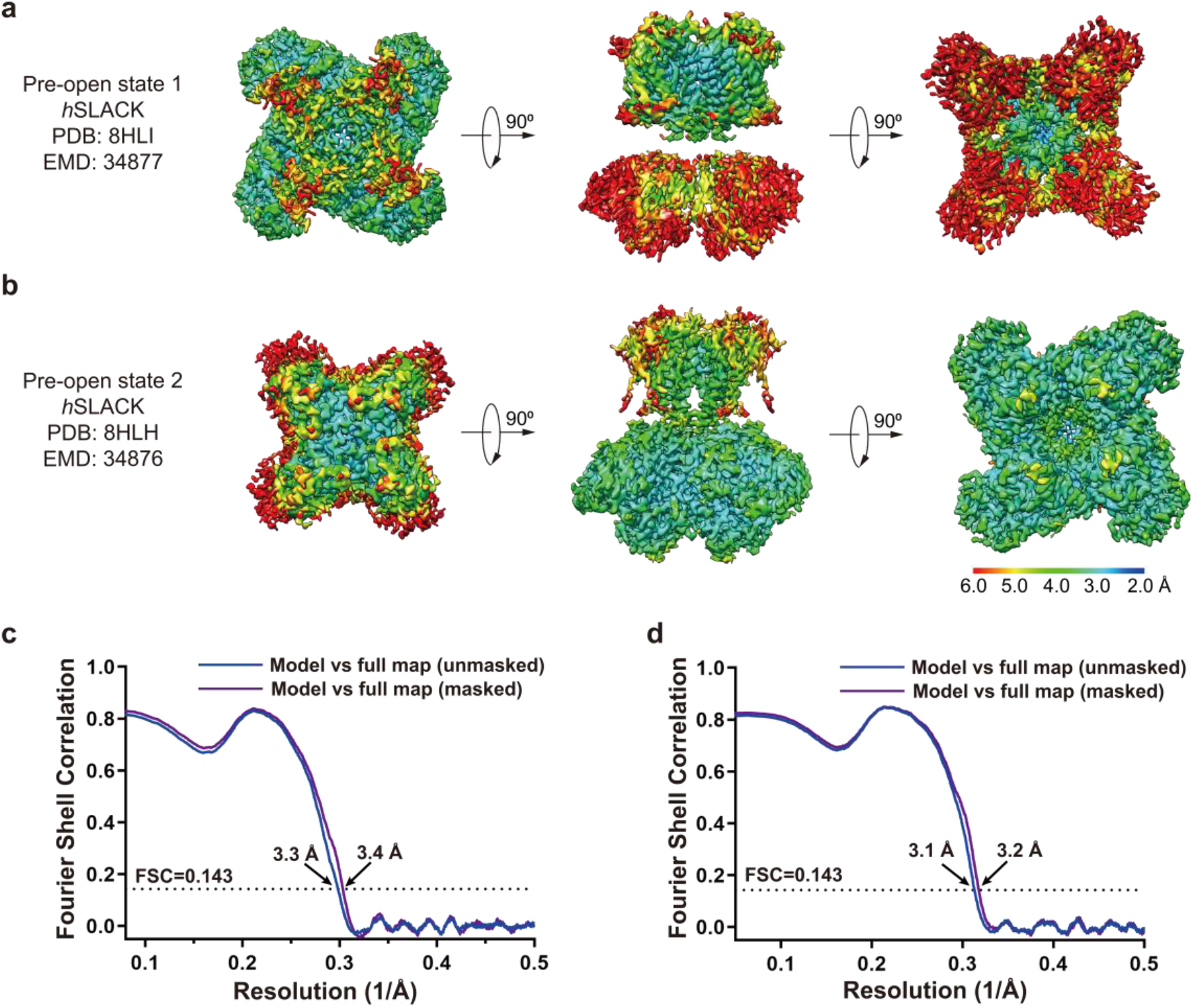
EM density of hSlack in pre-open state 1 and pre-open state 2. **a**, EM density of transmembrane helices of hSlack in pre-open state 1. **b**, EM density of hSlack intracellular region in pre-open state 1.

**Table S1.** Cryo-EM data collection, refinement, and validation statistics.

|  | Pre-open state 1 | Pre-open state 2 |
| --- | --- | --- |
|  | *h*SLACK | *h*SLACK |
|  | (PDB 8HLI) | (PDB 8HLH) |
|  | (EMD 34877) | (EMD 34876) |
| **Data collection and processing** |  |  |
| Microscope | FEI Titan Krios | FEI Titan Krios |
| Magnification | 105,000 | 105,000 |
| Voltage (KV) | 300 | 300 |
| Detector | Gatan K3 | Gatan K3 |
| Electron exposure (e<sup>-</sup>/ Å<sup>2</sup>) | 60 | 60 |
| Defocus range (μm) | -1.4 to -1.9 | -1.4 to -1.9 |
| Pixel size (Å) | 0.849 | 0.849 |
| Symmetry imposed | C4 | C4 |
| Initial particle images (no.) | 1,350,749 | 1,350,749 |
| Final particle images (no.) | 94,340 | 113,598 |
| Map resolution (Å) | 3.34 | 3.18 |
| FSC threshold | 0.143 | 0.143 |
| **Refinement** |  |  |
| Initial model used (PDB code) | 5U70 | 5U70 |
| Model resolution (Å) | 3.3 | 3.2 |
| FSC threshold | 0.143 | 0.143 |
| Map sharpening *B* factor (Å<sup>2</sup>) | 93.1 | 94.9 |
| Model composition |  |  |
| Non-hydrogen atoms | 29321 | 29355 |
| Protein residues | 3616 | 3616 |
| Water | 4 | 8 |
| Ion | 9 | 39 |
| Lipid | 4 | 4 |
| B factors |  |  |
| Protein | 47.77 | 43.06 |
| Ligand | 31.70 | 29.29 |
| Water | 40.75 | 19.37 |
| R.m.s. deviations |  |  |
| Bond lengths (Å) | 0.007 | 0.005 |
| Bond angles (°) | 1.039 | 0.893 |
| Valiation |  |  |
| MolProbity score | 2.37 | 2.22 |
| Clash score | 16.25 | 12.08 |
| Poor rotamers (%) | 0.00 | 0.12 |
| Ramachandran plot |  |  |
| Favored (%) | 84.91 | 86.83 |
| Allowed (%) | 14.20 | 12.28 |
| Disallowed (%) | 0.89 | 0.89 |

## Acknowledgments

We would like to thank the Cryo-EM Facility of Westlake University for providing cryo-EM and computation support. This work was supported by Westlake Laboratory (Westlake Laboratory of Life Sciences and Biomedicine) and an Institutional Startup Grant from the Westlake Education Foundation to D.P. We also would like to thank all the Cell fate control lab members for their support.

## Author contributions

M.Z. and D.P conceived the project. Y.S. and M.Z. designed the experiments. Y.S. prepared the cryo-EM sample and collected cryo-EM data. M.Z. performed image processing and analyzed EM data. Y.S. and M.Z. performed electrophysiology experiments and wrote the manuscript draft. All authors contributed to the manuscript preparation.

## Reporting summary

Further information on research design is available in the Nature Research Reporting Summary linked to this paper.

## Data availability

The coordinate files and cryo-EM maps of the pre-open state 1 and pre-open state 2 hSlack have been deposited at Protein Data Bank (Electron Microscopy Data Bank (EMDB)) under accession codes 8HLI (EMD-34877) and 8HLH (EMD-34876), respectively.

